# Spider viscid silk sticks better to superhydrophobic surfaces

**DOI:** 10.1101/2020.05.15.098194

**Authors:** Angela M. Alicea-Serrano, K Zin Htut, Alissa J. Coonfield, Katherine Karkosiak, Ali Dhinojwala, Todd A. Blackledge

**Affiliations:** Department of Biology and Integrated Bioscience, University of Akron, Ohio; Department of Polymer Science, University of Akron, Ohio

## Abstract

In a likely coevolutionary arms race, insects evolved a variety of counter strategies to avoid capture by spider webs, while spiders’ evolved innovations web structure and especially their adhesive silks. For instance, insects’ cuticles employ a variety of potential anti-adhesion mechanisms such as the detachable scales of moths and surface waxes and superhydrophobic structures that might resist spreading of glues. In contrast, the viscid capture threads of most spider orb webs are covered with aggregate glue droplets that absorb atmospheric water, tuning glue viscosity to balance the competing demands of surface spreading versus maintaining strong bulk cohesion. Here, we test the hypothesis that superhydrophobicity in insects acts as an anti-adhesion defense against spider silk. We used lotus leaves as a model substrate because its superhydrophobicity outperforms most known insect surfaces. The adhesion of spider capture silk from the web of *Larinioides cornutus* was studied against three substrates: raw lotus leaves, oxygen plasma treated lotus leaves (hydrophilic lotus), and smooth glass, differing in roughness and chemistry. We found that spider capture silk sticks better to the superhydrophobic lotus than to other surfaces. Both chemistry and physical properties of the leaves contribute to higher adhesion, as raw lotus showed a mean increase in adhesion of 74 % compared to glass, while the similar surface roughness of the hydrophilic lotus increased adhesion by 64 % compared to glass. Thus, evolving a hydrophobic cuticle is unlikely to be a defensive trait used to mitigate the effectiveness of spider webs.

## INTRODUCTION

Spiders are diverse and important predators of insects in nearly every terrestrial ecosystem on earth (Garrison et al., 2016; Vollrath and Selden, 2007). Spiders in the superfamily Araneoidea are among the most successful predators of flying insects due in part to their use of a wet sticky silk, known as viscid silk (Blackledge et al., 2009; Bond and Opell, 1998; Coddington and Levi, 1991; Opell and Bond, 2001; Tarakanova and Buehler, 2012). Viscid silk is used by modern orb weaving spiders to build the capture spiral of their web, which functions to retain prey hitting the web until the spider is able to subdue it (e.g. Blackledge and Zevenbergen 2006). Viscid silk is made of an axial fiber of flagelliform silk coated by evenly spaced glue droplets of adhesive silk from the aggregate gland, both of which stretch and deform during pull-off from prey by implementing a suspension bridge mechanism that recruits glue droplets to simultaneously transfer forces from droplet-to-droplet and from droplet-to-axial fiber (Opell and Hendricks 2007; Liao et al. 2015). Glue in viscid silk is made of glycoproteins, lipids, and hygroscopic low molecular mass compounds (LMMC) (Choresh et al., 2009; Jain et al., 2018; Townley and Tillinghast, 2013; Townley et al., 1991; Vollrath and Edmonds, 1989; Vollrath and Tillinghast, 1991; Vollrath et al., 1990). The water-soluble LMMC’s are humidity-responsive, and aid in plasticizing the glycoproteins –which serve as the adhesive component and are not soluble in water (Opell et al., 2011; Opell et al., 2013; Sahni et al., 2010; Sahni et al., 2011; Sahni et al., 2014)– and also remove interfacial water (Singla et al., 2018). Variation in composition of LMMC helps to tune glue hygroscopicity to achieve an “ideal” viscosity for a spider’s foraging environment by allowing the glue to spread enough to cover a high surface area but without risking premature cohesive failure from the thread (Amarpuri et al., 2015; Amarpuri et al., 2017; Jain et al., 2018). While the mechanism of glue adhesion is very well understood on smooth hydrophilic substrates, particularly glass, glue behavior on more realistic rough and hydrophobic substrates has not been invesstigated in detail.

In a possible coevolutionary arms race, insects evolved a variety of counter strategies to avoid capture by orb weaving spiders (Blackledge et al., 2009; Penney, 2004). These counter strategies include escape behaviors, sacrificial scales of moths and butterflies, and potentially the microstructure and chemistry of the chitin exoskeleton itself. Insect cuticles are multilayered composites of chitin and proteins and many cuticles are hydrophobic due to specialized protuberant structures or waxes present in the epicuticle (e.g. Watson et al. 2011). Cuticular waxes play a key role not only in preventing desiccation (Bott et al., 2017) but also hypothesized to function in several other ways, such as facilitating mating success (Chung and Carroll, 2015). Superhydrophobicity in insect cuticles may function in anti-fouling, anti-fogging, or anti-reflectance (e. g. Bott et al. 2017). Anti-wetting is also hypothesized to function as protection against spider webs by preventing effective spreading of adhesive glue droplets (Masters and Eisner, 1990; Watson et al., 2011). Some moth-specialist spiders in the subfamily Cyrtarachninae use large, over-lubricated glue droplets for quick spreading underneath superhydrophobic cuticles (Cartan and Miyashita, 2000; Diaz et al., 2018; Diaz et al., 2020) and cribellate spiders sometimes leverage capillary forces of epicuticular waxes of insects along the adhesive nanofibers to enhance adhesion (Bott et al., 2017). Most ecribellate orb weaving spiders, however, are prey generalists whose water-based glue is maximally adhesive when it balances the contributions of interfacial spreading and bulk cohesion during pull-off (Amarpuri et al., 2015). Thus, poor wetting of superhydrophobic cuticles could provide a very effective defense strategy that has been over-looked by most research on viscid silk function which investigates adhesion primarily on smooth, hydrophilic glass surfaces.

Insect cuticles vary widely in hydrophobicity—from cockroaches with a contact angle below 90° to mayflies, lacewing flies, butterflies, and moths with a contact angle ranging well above 130° (Wagner et al., 1996). Because much of this variation in hydrophobicity is confounded with macroscale features such as setae and sacrificial scales that can strongly influence adhesion, we chose to explore the effects of hydrophobicity using a different system – the lotus leaf. The lotus leaf ranks as one of the most super-hydrophobic surfaces in nature with a water contact angle of around 150°, giving it self-cleaning properties that are of great interest for biomimicry. Lotus leaf superhydrophobicity results from a hierarchical structure of randomly oriented hydrophobic wax nano-tubules on the top of convex cellular micro-papillae (Figure 1) (Barthlott and Neinhuis, 1997). However, the leaves are otherwise smooth, lacking hairs and other potentially entangling macro-structure, making them a good comparison to the commonly used glass surfaces used in various studies on spider silk adhesion. Here, we test the effects of substrate hydrophobicity *per se* on the adhesion of the spider silk, to test the hypothesis that superhydrophobicity can be utilized by insects to mitigate the effectiveness of spider viscid silk. Results from this project will help in the development of a more robust understanding of the ecology of orb weaving spiders and their success at catching flying prey, as well as the advancement of technology to create biomimetic adhesives.

**Figure 1.**
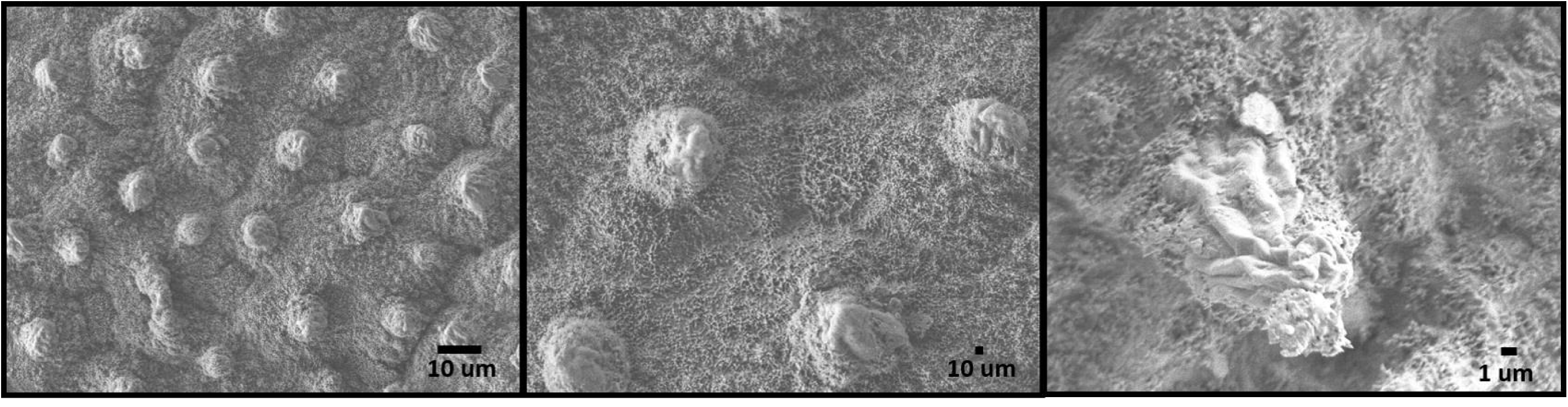
Lotus leaf is classic example of a superhydrophobic surface. This property is possible due to its hierarchical structure of micro papillae and nano wax tubules (not visible).

## METHODS

### Spider maintenance and silk collection

*Larinioides cornutus* (Araneae: Araneidae) orb weaving spiders were collected from two locations near Bath Township in Akron, Ohio, USA. Spiders were housed in cages (37cm width x 38 cm height x 11.5 cm depth) and kept at the University of Akron to build webs. Spiders were fed a cricket diet after every web collection (up to three times per week), and sprayed with water every afternoon before nighttime, the time at which *L. cornutus* normally builds. Single strands of capture silk were collected from interior rows in the bottom side of the web, by securing it across 12.58 mm cardboard frames using Elmer’s glue (©Borden Inc.) and then cutting using a soldering iron. Samples were stored in a dark sealed box, avoiding exposure to dust and UV light for up to 34 days until testing.

### Substrate preparation and surface energy testing

To test the adhesive properties of capture silk on a superhydrophobic substrate, lotus (*Nelumbo nucifera*) leaves were used as a substrate. Lotus leaves are notorious for their low surface energy and self-cleaning properties. Because of their hierarchical surface made of micro papillae and nano wax tubules, water cannot spread on its surface. To decouple the effect of surface chemistry from the surface topography, lotus leaves were plasma treated with oxygen for 5 minutes (Harrick Plasma, PDC-32G), making the leaves superhydrophilic by depositing a layer of OH ions at the outermost surface of the leaf. Capture silk was also tested on smooth hydrophilic glass as a control. Water contact angles were measured on both hydrophilic and raw lotus leaves with Kruss Dynamic Shape Analyzer by depositing 10 μL droplets on these surfaces to provide insights into the surface energy of these substrates. Six fresh leaves from four different plants were used and three droplets for each leaf were tested for determining average water contact angles. The resulting images were analyzed using contact angle analysis plug-in with image J software (Schneider et al., 2012).

### Adhesion testing

A total of 120 threads from 40 webs were tested across all three substrates (n = 40 each for raw lotus leaves, hydrophilic lotus leaves, and glass) so that any variation among webs in stickiness was evenly distributed across treatments. Substrates were made from the same leaves used in contact angle measurements, by adhering them to tick cardboard using double sided tape and cutting them into rectangles. All substrates had the same width (5 mm) to control for the length of capture silk coming in contact. Adhesive properties were tested using the Nano Bionix® Tensile tester (MTS Nano Instrument Innovation Center) by bringing individual threads in contact with the substrate and measuring pull-off forces. Threads were brought in contact with the substrate at a constant speed of 0.1 mm/s until a 15 nN preload, held in place for six seconds, and then pulled off until detachment using the same speed of 0.1 mm/s. Each thread was tested once on a discrete part of the substrate. All tests were done at 60 ± 5 % relative humidity, the condition at which the glue of *L. cornutus* shows maximum adhesion on glass (Amarpuri et al., 2015). Humidity was controlled inside a humidity chamber using nitrogen gas and de-ionized water. Work of adhesion, or energy it takes to unstick from the substrate, was calculated as the area under the load-extension curve. Peak load during pull-off and total extension of the capture silk were also measured. A one-way analysis of variance (ANOVA) was used, with an alpha level of 0.05, to determine if the average per web work of adhesion, peak force of adhesion and extension were different for each substrate. Tukey HSD was performed *post hoc* to determine pairwise differences among groups per variable. All statistical testing and graphing was performed using the JMP® Pro 14 software (©SAS Institute Inc.).

### Imaging

A scanning electron microscope (SEM) (Model JEOL-7401 Japan Electron Optics Laboratory (JEOL)) was used to visualize (1) the morphology of the lotus, (2) the glue sticking behavior on the lotus leaf, and (3) the substrates after adhesion tests. Samples for 1 and 2 were observed in their native state without any sputter coating, in high vacuum operating at 1-3 kV, and were also visualized using a Keyence© VHX-7000 digital microscope (Keyence Corporation of America). Two leaves were used to observe the morphology of lotus before and after the oxygen plasma treatment. For 3, all substrates used in adhesion testing were visualized after testing to see if any glue residue was left behind. Samples were sputter coated for 45 seconds with gold, and observed in high vacuum operating at 1-3 kV. Any cohesive failure observed was recorded by counting the number of threads leaving visible glue residue (Figure 4 b1 and c1). A percentage of cohesive failure was calculated for each substrate by dividing the number of threads with cohesive failure by the total number of threads tested in that substrate.

**Figure 2.**
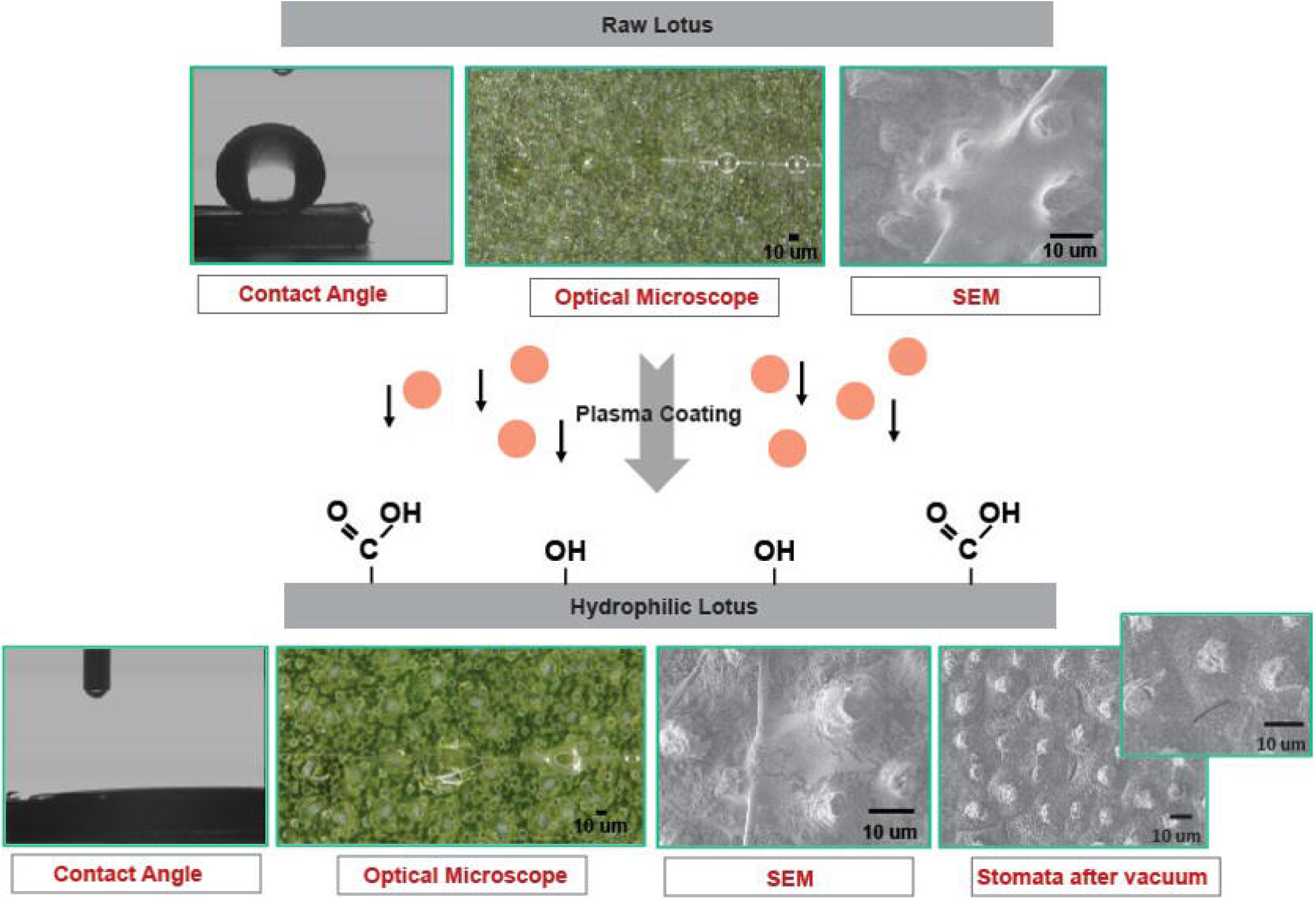
Diagram showing the process of oxygen plasma-treatment. Plasma-treatment results in oxidation of non-polar methyl and methylene groups of waxes to polar carboxylic and hydroxyl groups using oxygen ions available in air plasma changing the chemistry from a non-wetting to a complete wetting state. It dramatically increases the ability of water to wet the leaf surface.

**Figure 3.**
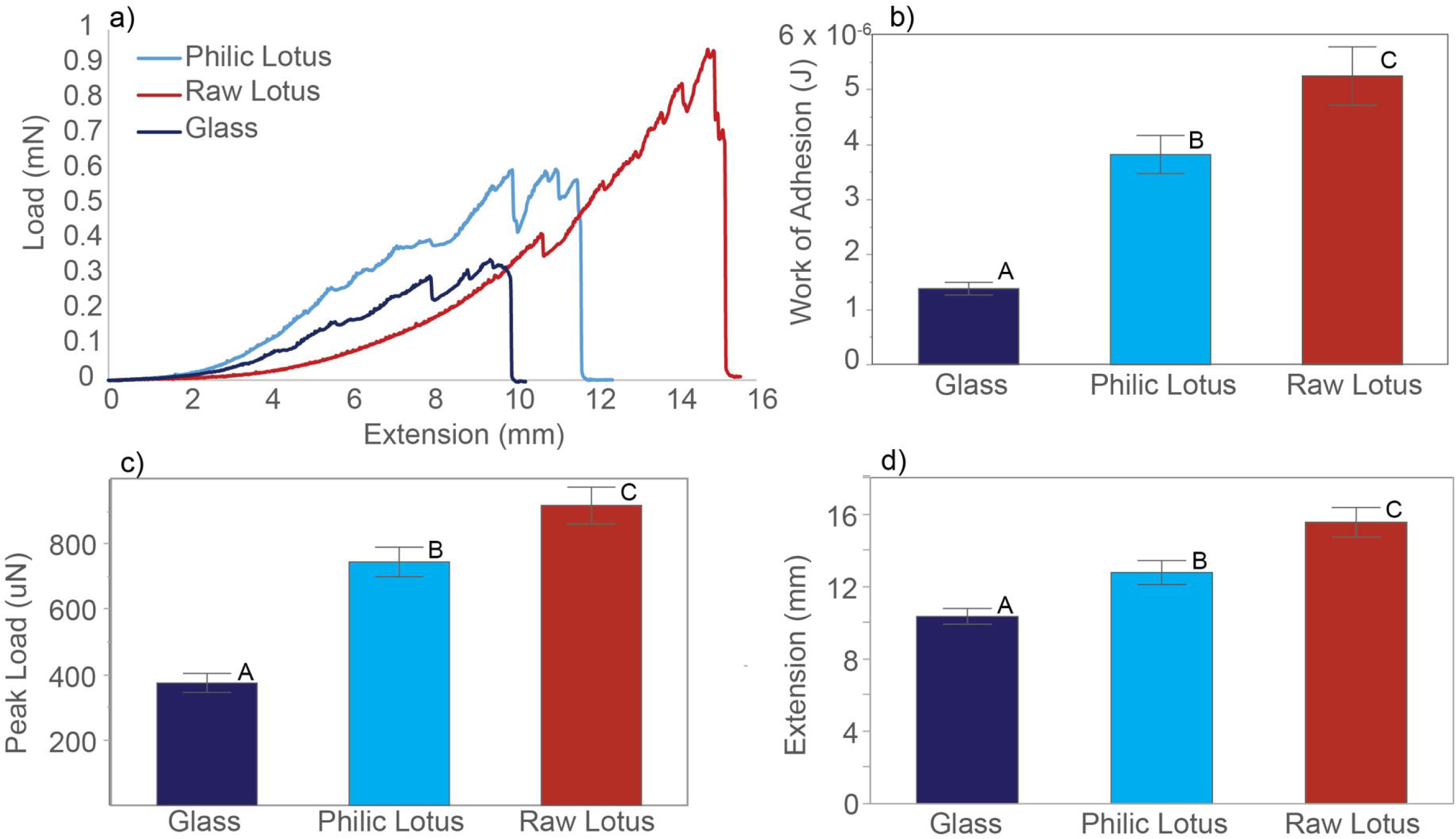
Viscid capture silk pull-off testing results for three substrates: (red) raw lotus, (light blue) hydrophilic lotus and (dark blue) glass. a) Load-extension curves representative of the average performance of capture silk on each of the substrates. b) Work of adhesion results showed glass had the lowest adhesion while the raw lotus had the higher adhesion. c, d) Work of adhesion was driven by both force of adhesion and extension of the capture silk as both showed the same trend as that observed in the work of adhesion.

**Figure 4.**
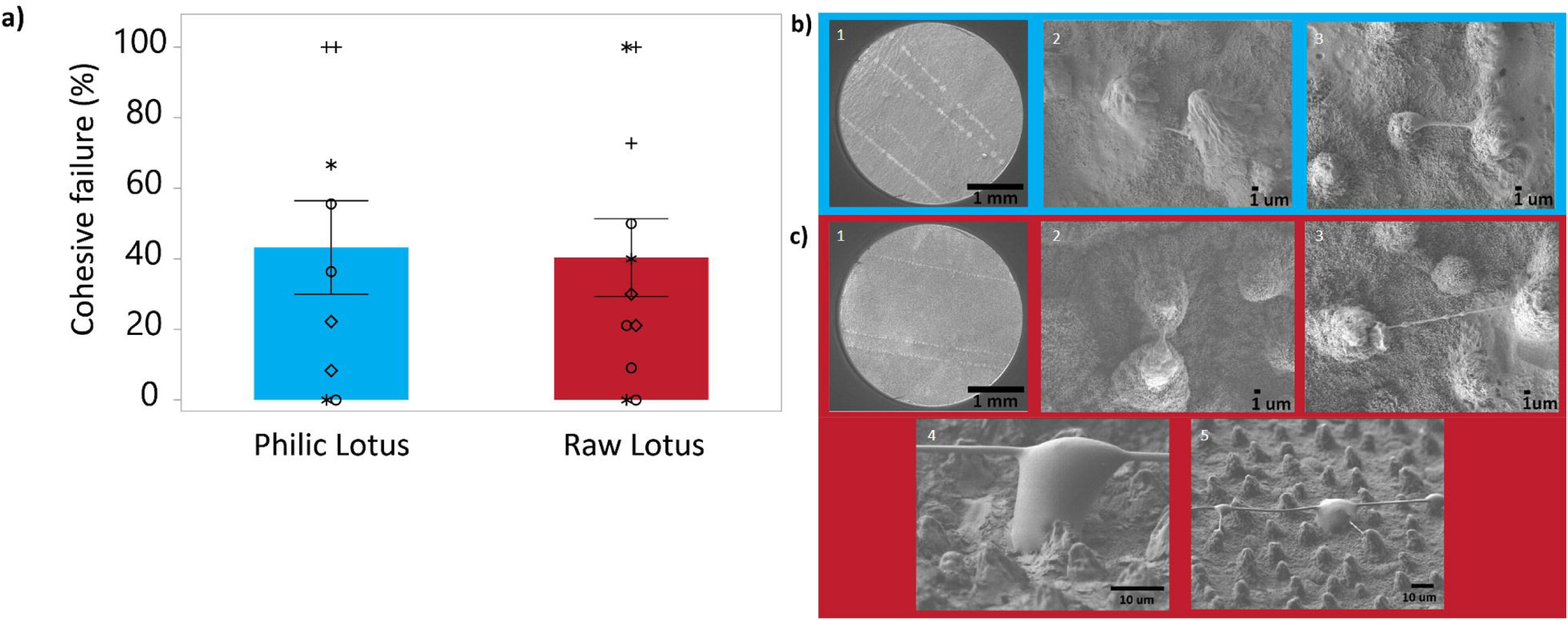
a) Percentage of cohesive failure of glue from capture silk observed on hydrophilic and raw lotus was the same for both treatments. Multiple papillae are cover by a single droplet of the capture silk increasing the surface area of contact and glue tendrils formed on the peaks of papillae. Tendrils observed on (b) hydrophilic lotus were scarce and only happened in really closely together papillae causing them to be short. Tendril formation was almost exclusively found in (c) raw lotus.

## RESULTS

### Surface energy

Lotus leaves have a low surface energy and self-cleaning properties due to a hierarchical surface made of micro papillae and nano wax tubules, on which water cannot spread. To decouple these two confounded factors (surface chemistry and topography), we oxygen plasma treated lotus leaves making them hydrophilic. Both hydrophilic substrates (hydrophilic lotus and glass) showed contact angles of zero degrees (°) while, raw lotus leaf had an average contact angle of 143.6 ± 1.2 ° (Figure 2). Visually, the stomata (pores found in leaves for the exchange of gas and water) in hydrophilic lotus leaves looked conspicuous under SEM pictures due to some process of plasma treatment, while they were unnoticeable for raw lotus (Figure 2). No other visual differences between the two lotus treatments were observed. Therefore, we defined our three substrates as smooth / hydrophilic (glass), rough / hydrophilic (plasma treated lotus) and rough / hydrophobic (raw lotus).

### Adhesion

We tested the adhesive properties of capture silk from *Larinioides cornutus*’ webs on raw (rough/ hydrophobic) and plasma treated (rough/ hydrophilic) lotus leaves and glass (smooth/ hydrophilic) substrates, which have varying roughness and surface chemistry. We found that overall capture silk outperforms on hydrophobic lotus than the other two substrates. Adhesion was higher on raw lotus leaf with a mean work of 5.25 × 10^−6^ ± 5.3 ×10^−7^ J, 1.5 times higher than the adhesion of capture silk on hydrophilic lotus (3.82 × 10^−6^ ± 3.48 ×10^−7^ J) and ∼4 times higher than work of glue pulling-off from glass (1.38 × 10^−6^ ± 1.15 ×10^−7^ J) (F (2,177) = 27.5958, *p* < 0.0001; Figure 3 a, b). The differences in the work of adhesion for the three substrates were driven by both the force during pull-off and the extension of capture silk. Hydrophobic lotus had both a significantly higher pull off force and a higher extension (918 ± 56.2 μN; 15.5 ± 0.8 mm), followed by hydrophilic lotus (745 ± 45.1 μN; 12.8 ± 0.7 mm) and glass (375 ± 28.8 μN; 10.3 ± 0.4 mm) (F (2, 117) = 38.3490, *p* < 0.0001; F (2, 117) = 15.6560, *p* < 0.0001; Figure 3 a, c, d).

After substrates were used to test adhesion, we examined them to see if glue residue was left behind. We found that about 40 % of capture silks tested on raw and hydrophilic lotus left glue behind due to cohesive failure inside the glue (Figure 3). Cohesive failure was observed to be equally abundant on both raw and hydrophilic lotus (Figure 3 a). An interesting behavior was observed for the capture silk glue as it interacted with the lotus rough surface, where glue tendrils formed at the peaks of papillae (Figure 3 b, c). This behavior was almost exclusively found on the residual glue on raw lotus leaves, while it was only observed in a few instances for hydrophilic lotus. Furthermore, tendrils observed on hydrophilic lotus only occurred in places where papillae were close together causing these tendrils to be very short and thick relative to the ones found on raw lotus.

## DISCUSSION

We tested the hypothesis that superhydrophobic surfaces could mitigate the effectiveness of spider capture threads by comparing spider silk adhesion on raw lotus leaves to glass and hydrophilic plasma-treated lotus leaves. We rejected the hypothesis because spider silk adhered most effectively to raw superhydrophobic lotus leaves compared to glass, the substrate mostly used to test glue stickiness. The ability of aggregate glue to effectively adhere to superhydrophobic substrates implies that insects have limited potential to evolve cuticular chemistries and micro-morphology as defenses against spider webs.

Variation in cuticular chemistry and microstructure plays a key role in how insects interact with their environment (Gullan and Cranston, 2014). For instance, specific cuticular microstructures can help insects thrive in aquatic, aerial, and arid environments (e.g. plastrons help with oxygen retention in aquatic environments; bristles, scales and microtrichia can decrease friction between air and cuticle to enhance aerodynamics; and leaf-like bristles aid in water retention useful in arid conditions) (Gorb, 2001). Other insects may have evolved cuticular features that function as anti-predator specializations against ubiquitous spider webs in the environment. For example, scales of butterfly/moths and, hairs in caddisfly and lacewing allow them to rapidly escape most orb webs by scarifying or shedding these layers (Eisner et al., 1964; Masters and Eisner, 1990; Nentwig, 1982). Indeed, in a coevolutionary arms race, insects and spiders potentially drove each other into diversification (Vollrath and Selden, 2007). However, our results argue that cuticular hydrophobicity and topography is an unlikely strategy to evolve as a general defense against gluey spider silks.

Unlike glue from other systems, like the one produced by the pupa of fruit flies, that successfully adhere to substrates with different roughness and chemistries (Borne et al., 2020), we found that glue adhere better to lotus leaf than smooth glass. Roughness is more important than chemistry for explaining the higher adhesion of capture silk on lotus leaf compared to glass (raw hydrophobic lotus: 74% vs hydrophilic lotus: 64 % change from glass). Roughness likely improves adhesion in two ways, increasing the total surface area for glue to adhere to in a given space and incorporating shear forces into adhesion. Previous work on the effect of macroscale cuticular features, such as hairs and setae, on adhesion also found that roughness could improve adhesion by increasing the total effective surface area of adhesion when setae were smaller than glue droplets but reduced adhesion when setae were much larger than glue droplets and therefore prevented effective contact (Opell and Schwend 2007). Capture silk used in our study, from webs of *Larinioides cornutus* have droplets that are larger than lotus papillae, and many of these papillae can be covered by a single droplet (Figure 2).

The 10 % increase in stickiness due to chemical differences between hydrophobic lotus and the plasma-treated hydrophilic lotus can be explained by overspreading of the glue (Amarpuri et al. 2017, 2015). Due to overspreading the contact line gets pinned and the stress is experienced by a small volume of glue, and this leads to an early cohesive failure. This was also reflected in lesser extension during retraction for plasma-treated hydrophilic compared to hydrophobic lotus. For hydrophobic lotus we also observed a novel behavior where tendrils or filaments of glue formed at the peaks of lotus papillae as glue pulled off from the substrate. These tendrils occurred almost exclusively on raw lotus. We believe that the retraction of glue on hydrophobic lotus is much smoother and gets pinned only at the tip of the papillae. This results in the glue getting drawn into thin filaments (Figure 4c, 5b) resulting in the formation of beads on a string (BOAS) morphology (Figure 4c), similar to the thinning behavior of saliva stretched between a thumb and forefinger (Schipper, Silletti, and Vingerhoeds 2007). The stretching of the filaments also results in higher extensibility. The tendril formation was not observed for hydrophilic lotus (observed in less than 1% of the strand tested) because the glue droplet has spread further and was pinned on locations other than the papillae. Thus, we think that during pull-off of viscid silk from the lotus substrate, large volume of the glue participates in resisting the pull off, resulting in higher extensibility (Figure 3 d), higher peak loads (Figure 3 c) and formation of filaments that are thinner and, in some cases, BOAS (Figure 4 c). In both cases, glue is left behind on the surfaces, indicating that a complete detachment of the glue does not take place on both raw and hydrophilic lotus. In future, imaging the glue droplet during spreading and detachment is necessary to confirm this proposed mechanism. Additionally, viscid glue may generate additional adhesive forces through physical interactions with surface lipids since superhydrophobicity of lotus is only for water and it may not be for glue droplets that contain LMMC or other lipid like compounds. However, testing this hypothesis is out of the scope of this work.

**Figure 5.**
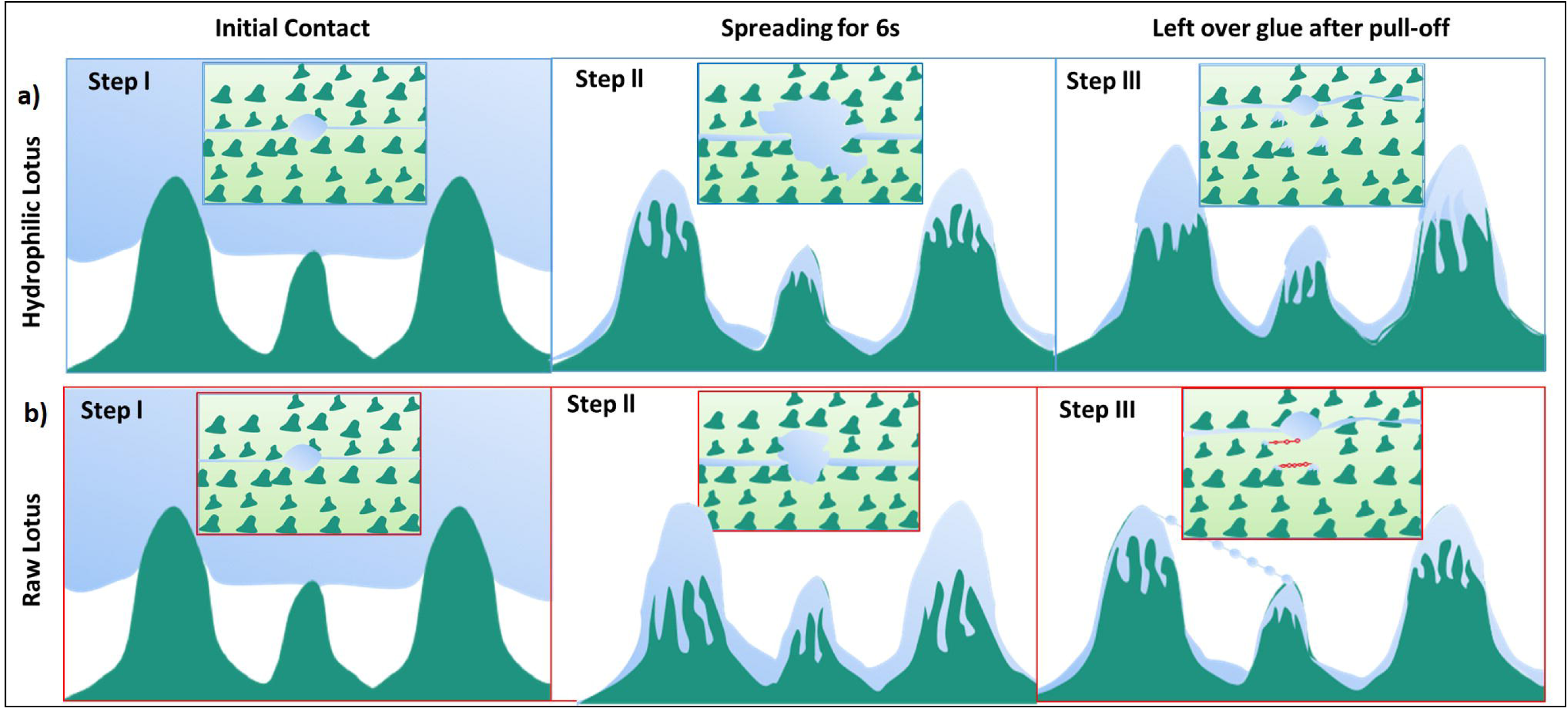
Hypothetical model for viscid glue pull-off from a) hydrophilic lotus and b) raw lotus. Droplets on hydrophilic lotus spread thinly, coating a large area but causing failure in the bulk at the periphery. Glue on raw lotus spreads ideally, allowing bulk cohesive forces to resist deformation at the periphery, while tendrils formed at peaks in papillae due to thinning.

Our study found that spider capture silk sticks better to the superhydrophobic surface of lotus. This finding is highly important for the understanding of the coevolution between insects and spider’s gluey silk. Although other factors, such as web architecture and both insect and spider behavior contribute to the spider’s success in trapping prey, our results show that evolving a superhydrophobic surface does not defend insects against spider webs, and capture silk. Humidity where adhesion is maximum may be a function of surface roughness and chemistry and therefore, understanding adhesion behavior of spider glue on heterogeneous surfaces is essential for developing tunable adhesives with multi-functional capabilities for biomedical, industrial, and robotic fields due to their unique and tunable viscoelastic properties.

## ACKNOWLEDGEMENTS

We would like to thank Kaleb Wells for his help in data collection, spider care and plants care, and for silk collection. We would also thank Saranshu Singla and Daniel Maksuta for reviewing and useful comments during the development of this study and the manuscript.

